# Transfer Learning Assessment of Data-driven Crystallisation Processes via Constrained Neural Ordinary Differential Equations

**DOI:** 10.1101/2025.09.22.677735

**Authors:** Daniele Pessina, Tony Tian, Oliver Watson, Jerry Y. Heng, Maria M. Papathanasiou

## Abstract

Modelling complex crystallisation processes remains challenging due to limited experimental datasets, high measurement noise, and the need for generalisability across varying operating conditions. Neural Ordinary Differential Equations (NODEs) and transfer learning (TL) offer promising tools to overcome these limitations by providing data-efficient, flexible, and transferable modelling frameworks. This work investigates the use of NODEs to model protein crystallisation dynamics under data-scarce conditions. A NODE trained on a data-rich source system successfully captures solute consumption and particle size dynamics, but when applied to data-sparse target systems, scratch-trained NODEs exhibit limited generalisation and unphysical behaviours. To address this, several TL strategies are evaluated, including layer freezing, parameter deviation penalisation, and system-embedding within the neural architecture. Results show that layer freezing and deviation penalty consistently improve knowledge transfer, while system-embedding offers robustness in noisy or undersampled datasets. In addition, physics-informed NODEs, constrained to enforce monotonic concentration decay and crystal growth, demonstrate greater stability under high noise and sparse measurement regimes, ensuring physically consistent predictions. Overall, the combination of constrained NODEs with appropriate TL strategies provides a robust framework for accurate, transferable modelling of crystallisation systems in low-data regimes.

## 1. Introduction

Process models are foundational tools in process design, optimisation, and control. *In-silico* simulations enable the exploration of process dynamics under varying conditions and reduce the need for expensive and time-intensive experimental campaigns. As such, they can accelerate development cycles and enhance decision-making in complex manufacturing and bioprocess systems.

Mechanistic models are typically constructed by combining mass, energy, and conservation laws with physically-informed or empirically-derived kinetic relationships, and offer high interpretability and grounding in physical theory. Despite the level of detailed often captured in the formulation, mechanistic models are not always accurate representations of real-world system. Structural simplifications are often necessary, whether due to limited experimental accessibility of key variables, significant uncertainty in parameter estimation, or practical considerations such as the need to reduce computational complexity. The formulation process itself can be labour-intensive, and regardless of the large effort the resulting models may fail to capture complex system behaviour with sufficient fidelity.

As an alternative, data-driven modelling paradigms have gained traction for their flexibility and scalability, and have been successfully applied for process design, optimisation, and control (Fahmi et al. 2012; Rio-Chanona et al. 2019; Ma et al. 2024; Lima et al. 2022; Benyahia et al. 2021; Michalopoulou et al. 2025). Process Analytical Technology (PAT) tools have enabled frequent, high-resolution measurement of key process variables, providing rich data streams, and motivate the exploration of machine learning (ML) and data-driven modelling strategies that do not rely on detailed mechanistic knowledge. Owing to their adaptable model structure they can, in principle, approximate system dynamics more accurately, especially in regimes where mechanistic insight is limited or unavailable. However, this comes at the cost of interpretability: while black-box models may yield better predictive accuracy, they lack the transparency and physical coherence offered by mechanistic approaches.

### 1.1. Neural Ordinary Differential Equations

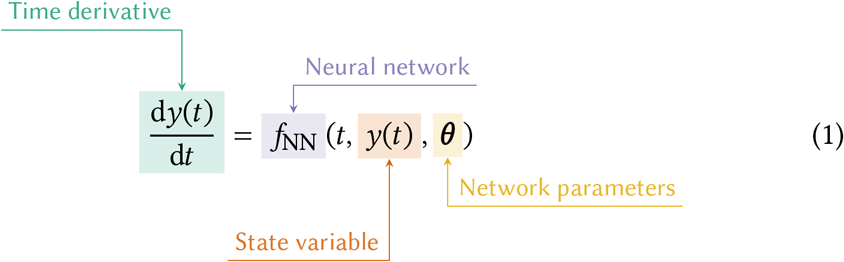

Neural ordinary differential equations (NODEs), introduced by Chen et al. (2019), are a promising ML modelling method in which a neural network (NN) is embedded within a differentiable ODE solver to approximate the derivatives of a system of ODEs (Eq. 1) (Kidger 2022). NODEs represent a recurrent neural network in which the time steps Δ*t* between hidden states *h*_*t*_ and *h*_*t*+1_ are reduced to zero and the formulation is then cast as the solution to a differential equation. Importantly, the NODE does not learn a single solution to the ODE, but rather approximates the *ODE flow*, the collection of ODE trajectories for all valid initial conditions and timepoints (Immler et al. 2016). For this reason, NODEs have good extrapolative capabilities and offer great potential towards improved process understanding. By decoupling the time discretisation from the ML model, NODEs are also compatible with irregular sampling times, missing data, and arbitrary residence times, an advantage over pure neural-network based data-driven modelling solutions (Luo et al. 2023).

Although the NODEs themselves remain black-box models, if their input features are mapped to relevant process variables (i.e temperature, pressure, concentration of a reactant), they can be used for enhanced interpretability and parameter estimation, as the NODEs give access to state gradient information (Bradley et al. 2024; Bradley et al. 2021). Kumar et al. (2024) directly embedded physical constraints in the NODE structure to predict stiff reaction kinetics, and achieved improved training efficiency. Fedorov et al. (2023) developed a kinetics-constrained NODE for improved training efficiency with small datasets. Yin et al. (2023) proposed a generalised NODE reactor model that achieved improved interpretability and data efficiency compared to other ML modelling methods. Chiu et al. (2025) used NODEs to model the growth of Chinese Hamster Ovary cell in a bioreactor, showing that NODEs could more accurately capture non-linear dynamics compared to a mechanistic and hybrid model. Luo et al. (2023) used NODEs for model-predictive control; by leveraging access to the derivative information and compatibility with irregular sampling from sub-sampling methods for noisy measurements, model-predictive control performance was improved compared to traditional methods. The NODEs have also been shown to remain robust with noisy measurements, which might come from in-line PAT (Goyal et al. 2023), and are highly effective with sparse datasets, which can occur due to PAT failures (Bradley et al. 2024).

### 1.2. Transfer Learning

Despite their data efficiency, NODEs and ML models usually rely on high-quality datasets (Thebelt et al. 2022) which should uniformly cover a wide range of process conditions, as low variance data sets lead to poor extrapolating capabilities (Bradley et al. 2022). Actions such as catalyst or reactor unit replacement, or a shift in operating conditions, can make trained ML models unable to accurately predict under this new domain. This aspect prevents the successful application of ML modelling methods on new processes that have not yet been extensively measured. However, processes do not exist in isolation, and there are often analogous systems that are, in fact, well characterised. Transfer learning (TL) can be used to leverage existing knowledge and data of a related system to accurately model the target system. There are many possible TL approaches, each of which has been successfully used to improve ML modelling of processes with limited data, and the success of each technique depends both on the modelled system and the structure of the source ML model (Rogers et al. 2022).

Broadly, TL approaches start with a *source* neural network trained on a high-quality dataset to obtain accurate predictions of the source domain, and a *target* domain for which there is not enough data to correctly train a new network. The network parameters are assumed to collectively represent the network’s learned ‘knowledge’ of the source domain. The simplest TL method, fine-tuning, involves initialising the target network parameters from those of the source network (rather than random initialisation), after which the network is trained on the target dataset. Other methods instead ‘freeze’ the source NN parameters, and only the remaining parameters are fine-tuned on the target dataset. Alternatively, deviations between source and target network parameters can be penalised during model training (Zhuang et al. 2021). However, selecting the best layers to freeze or penalise is unclear. Laptev et al. (2018) suggested that the first layers of a network are ‘feature’ layers, and the latter are ‘predictive’ layers, and feature layers should be frozen for effective knowledge transfer. In the work the authors iteratively froze the frontal layers of a recurrent NN, identifying an optimal number of transferred layers. Riezzo et al. (2025) modified the loss function of a NN embedded in a hybrid model of microalgal production to penalise deviations of the weights and bias parameters from the original network. Xiao et al. (2023) and Alhajeri et al. (2024) modified the existing recurrent NN by freezing the original network parameters, adding ‘adaptation’ layers to the network and progressively training the new network. Rogers et al. (2022) compared these latter two methods, showing that weight penalisation appears to lead to improved knowledge transfer across the two domains.

The layer-freezing and deviation penalty methods re-purpose a trained NODE to instead predict the dynamics of the target system. A limitation is that they only consider pairs of source and target systems, transferring knowledge from the former to the latter. If data from multiple target systems is available, it would be beneficial to attempt at embedding, within the model architecture itself, the concept of multiple systems. Sitapure et al. (2023) made use of the attention mechanism of a time-series-transformer model to enhance system-to-system transferability across crystallisation systems. The transformer model showed improved performance compared to an equivalent LSTM model when predicting an unseen system. This was regardless of the fact the transformer model had never been trained on data from this new system, as the attention heads were able to identify structural similarities between the new system and its’ training data.

### 1.3. Template-induced protein crystallisation systems

Protein crystallisation systems are an example of well-characterised systems with data that can be leveraged to adapt a data-driven model to a new, data-scarce target domain. Proteins have limited crystallisation conditions due to their size, large number of bonds and flexible structure. High API concentrations might be needed, exceeding those eluted upstream (Hekmat et al. 2017; McCue et al. 2023). Recent publications have explored the use of nucleation-promoting additives such as silica nanoparticles or dissolved amino-acids, known as templates, to facilitate crystallisation at gentler conditions (McCue et al. 2023; Chen et al. 2020a; Chen et al. 2020b; Link 2022). In previous work, template-induced protein crystallisation systems were investigated, and the templates’ effect on crystallisation kinetics was assessed through the use of a mechanistic population-balance model (Pessina et al. 2025). The templates act as nucleation promoters, appearing to lower the estimated crystal-solution interfacial energy compared to homogeneous (template-free) crystal nucleation and generally lowering the supersaturation-dependence of nucleation, and they offer great improved process intensification for the crystallisation of bio-macromolecular APIs. While *in-silico* simulations of template-induced crystallisation systems can be used to explore their effect on Critical Quality Attributes and the whole crystallisation operating space, templates have seen limited testing at larger processing scales (Chen et al. 2021). Poor data availability prevents wider application of data-driven modelling to crystallisation processes.

To address the modelling challenges introduced by scarce crystallisation datasets, this work assesses the performance of knowledge transfer techniques for NODEs of protein crystallisation systems. The moments-based population balance model and kinetic parameters from Pessina et al. (2025) are used for this case-study, and are also collected and explained in Appendix A. NODEs are trained on *in-silico*-generated concentration and average particle size profiles of four crystallisation systems. Layer freezing, network parameter deviation penalty and system-embedding methods are assessed in their effectiveness at transferring knowledge from a source to target domains.

## 2. Methodology

### 2.1. Case-study: Template-induced protein crystallisation models

*In-silico* simulations are used to generate the necessary data to train the NODEs and assess the knowledge-transfer techniques. The sets of kinetic parameters describe 4 batch and iso-thermal anti-solvent lysozyme crystallisation processes: a homogeneous crystallisation system **S**, and 3 template-induced crystallisation systems in which porous silica nanoparticles with chemically-functionalised surfaces (hydroxyl **T**_1_, carboxyl **T**_2_ and butyl **T**_3_) are added to the crystallisation solution to promote crystal nucleation. The solute concentration *c*(*t*) and volume-weighted average particle size *d*_43_(*t*) are assumed to be the only two experimentally-available variables of the process. The complete ODE model (Appendix A) is solved for different initial concentrations (found in Table 1) and zero initial crystals and average particle size to generate training and testing trajectories for the NODEs. Simulated trajectories are reported in Figure A.17. The initial concentrations used to generate data for **T**_1-3_ are purposefully restricted to simulate low-variance datasets which only cover a narrow range of possible crystallisation conditions. The number of timepoints and the amount of added gaussian noise is later varied to assess the NODEs’ robustness. Mean absolute errors are calculated against reference simulations with 20 timepoints and zero added noise.

**Table 1:** Summary of the initial concentrations used to generate training and testing data.

| Initial concentrations for training |  |
| --- | --- |
| **Source** | 15.5, 16.5, 19 [mg/ml] |
| **T1-3** | 16.5, 17 [mg/ml] |
| Initial concentrations for testing |  |
| **Source, T1-3** | 14, 17.7, 20 [mg/ml] |

### 2.2. Neural ODE training

Before any neural network is trained, the generated data is pre-processed. The state variables are rescaled to the [-1, 1] range, and the time domain is normalised to [0, 1] to improve numerical conditioning and facilitate gradient-based optimisation. Neural networks are constructed using the equinox JAX package and embedded within a differentiable ODE solver from diffrax (Kidger et al. 2021; Kidger 2022). The NODEs are first trained on only the first 33% of each trajectory’s time points before training on the full dataset to prevent convergence on local minima. Each network’s hyperparameters are collected in Table 2 and 3. The NODEs used are *augmented* NODEs, in which additional dimensions with initial condition equal to zero are appended to the input dimensions of the network (Dupont et al. 2019). This allows the networks to make use of the additional dimensions to approximate the ODE flows at a reduced computational expenditure. The NODEs presented all use the swish activation function, are trained using the AdamW optimiser with weight decay regularisation and the mean squared error is minimised (Eq. 2). The AdamW optimiser and its weight decay regularisation is opted for due to improved generalisation over the Adam optimiser and lower risk of overfitting (Loshchilov et al. 2019).

**Table 2:**
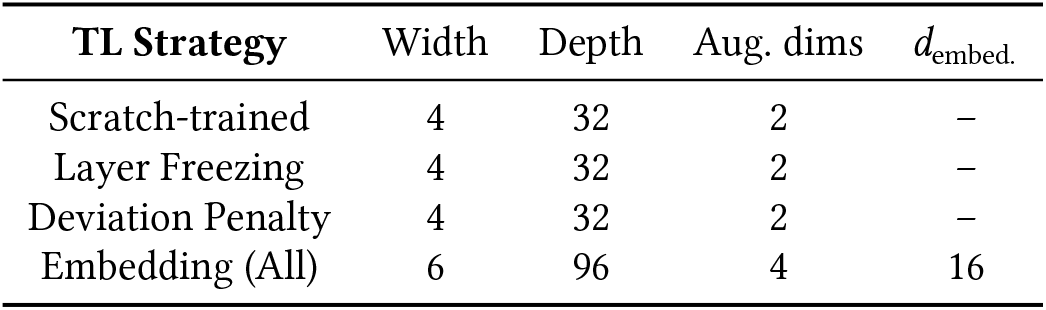
Neural network structural hyperparameters.

**Table 3:** Neural network training hyperparameters.

| TL Strategy | LR<sub>1</sub> | LR<sub>2</sub> | Ep<sub>1</sub> | Ep<sub>2</sub> | LR<sub>3</sub> | LR<sub>4</sub> | Ep<sub>3</sub> | Ep<sub>4</sub> | Dev. Pen. |
| --- | --- | --- | --- | --- | --- | --- | --- | --- | --- |
| Scratch-trained | $2 \times 10^{-4}$ | $4 \times 10^{-4}$ | 600 | 600 | – | – | – | – | – |
| Layer Freezing | $2 \times 10^{-4}$ | $4 \times 10^{-4}$ | 600 | 600 | $6 \times 10^{-4}$ | $6 \times 10^{-4}$ | 600 | 600 | – |
| Deviation Penalty | $2 \times 10^{-4}$ | $4 \times 10^{-4}$ | 600 | 600 | $6 \times 10^{-4}$ | $6 \times 10^{-4}$ | 600 | 600 | $1 \times 10^{-4}$ |
| Embedding (All) | $4 \times 10^{-4}$ | $6 \times 10^{-4}$ | 600 | 600 | – | – | – | – | – |

Gradients are calculated through reverse-mode autodifferentiation and the adjoint sensitivity method from diffrax. The Tsit5 solver is used for forward-in-time integration (Tsitouras 2011). A workflow of the NODE structure is shown in Figure 1.

**Figure 1:**
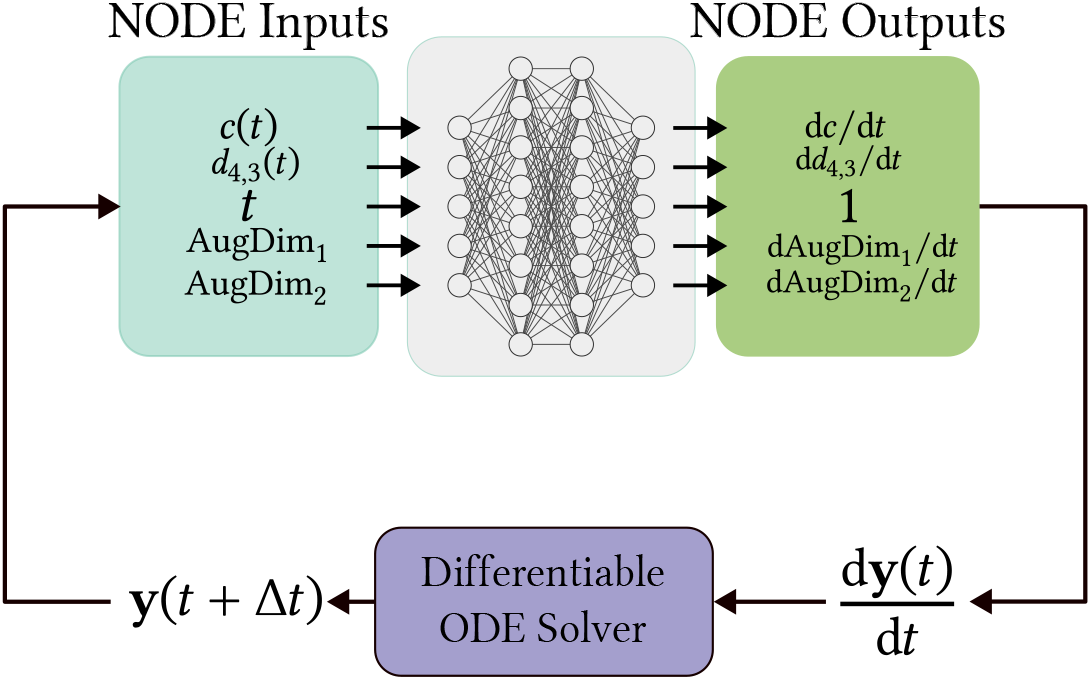
Neural ODE structure

#### 2.2.1. Constrained Neural ODEs

The NODE structure can also be constrained to enforce known physical restrictions of the system, such as monotonicity of specific process variables. For example, in the context of batch isothermal crystallisation systems, the solute concentration monotonically decreases, while the average particle size will monotonically increase. In this work the NODE outputs are constrained by applying a softmax function to the outputs during training.

### 2.3. Transfer Learning methods

Several different TL methods are assessed in this work. A scratch-trained NODE (NODE_**S**_) is first trained from the **S** data. TL methods are then applied, continuing the training on the **T**_i_ data.

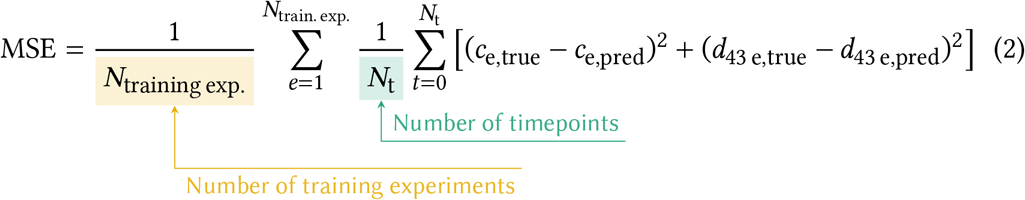

#### 2.3.1. Layer Freezing

With layer freezing methods, a subset of the weights and biases from NODE_**S**_ are fixed and become un-trainable. 5 layer freezing strategies are assessed by freezing layers L1, L1+2, L1+2+3, L3+4 and L4.

#### 2.3.2. Parameter deviation penalty

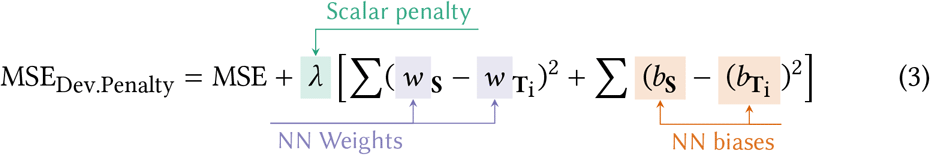

In the deviation penalty method, the mean squared difference between weights and biases of NODE_**S**_ and 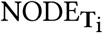 is summed, multiplied by a scalar parameter *λ*, and added to the MSE loss. The *λ* parameter tunes knowledge transfer (high *λ*) and fitting performance to the target dataset (low *λ*). In the implementation from Riezzo et al. (2025), the deviation of all network parameters is calculated. In this work specific layer selection for added penalty term is instead assessed.

#### 2.3.3. System-embedded NODEs

A different approach involves directly embedding knowledge of multiple domains within the model architecture, and conditioning the NODE with a zero-derivative input feature that is unique to this system. In this case, an initial raw look-up table is formed by stacking and normalising 4 one-hot system embedding vectors into a 4×4 matrix, with each row corresponding to the encoding of a different system. Depending on the selected system, the corresponding row is passed to a linear network layer, increasing its’ dimensions to a richer embedding of dimensions *d*_embed._ with a weighing matrix and bias parameter. Finally, this latent, higher-dimensional transformation returns a vector of length *d*_embed._ which is appended to the input features of the NODE. All of these parameters, including the original embeddings, are trainable and adapt to the modelled systems and their training data.

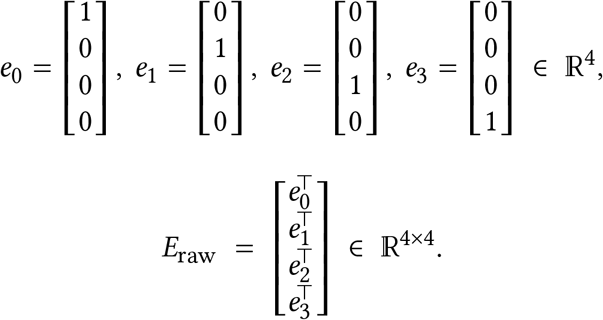

## 3. Results and Discussion

A scratch-trained, unconstrained NODE is first trained on the **S** dataset to be used for knowledge transfer. The training and testing predictions are shown in Figure 2. For each experiment in this and further figures, and in the error assessments, an ensemble of 5 NODEs is trained using different random seeds to account for variability in initialization and optimization, and the figures report the mean predicted trajectories and standard deviations. The NODE demonstrates good fit against the training set, both for the solute concentration and the *d*_43_ profile, indicating satisfactory learning of the underlying dynamics. On the test set, the NODE generalizes well for all three experiments though slightly poorer performance is observed when predicting the experiment at the lowest initial concentration. While the *d*_43_ trajectory remains well-predicted, a small unphysical increase in solute concentration is observed. Since the modelled crystallisation systems are iso-thermal and batch, the concentration should monotonically decrease while the average particle size should monotonically increase. Given that the lower-concentration trajectory lies furthest from the training data, the NODE may be extrapolating based on biased learned dynamics. The insufficient diversity in the training experiments and their initial conditions likely causes the model to prioritize high-density regions in ODE flow space, resulting in unstable and incorrect predictions. Figure 3 reports the prediction profiles for a constrained NODE trained on the **S** data, enforcing negative concentration gradients and positive *d*_43_ gradients. While the unphysical concentration increase is corrected, constraining the NODE leads to poorer particle size predictions for that experiment, with *d*_4,3_ growth excessively slowing towards the end of the batch.

**Figure 2:**
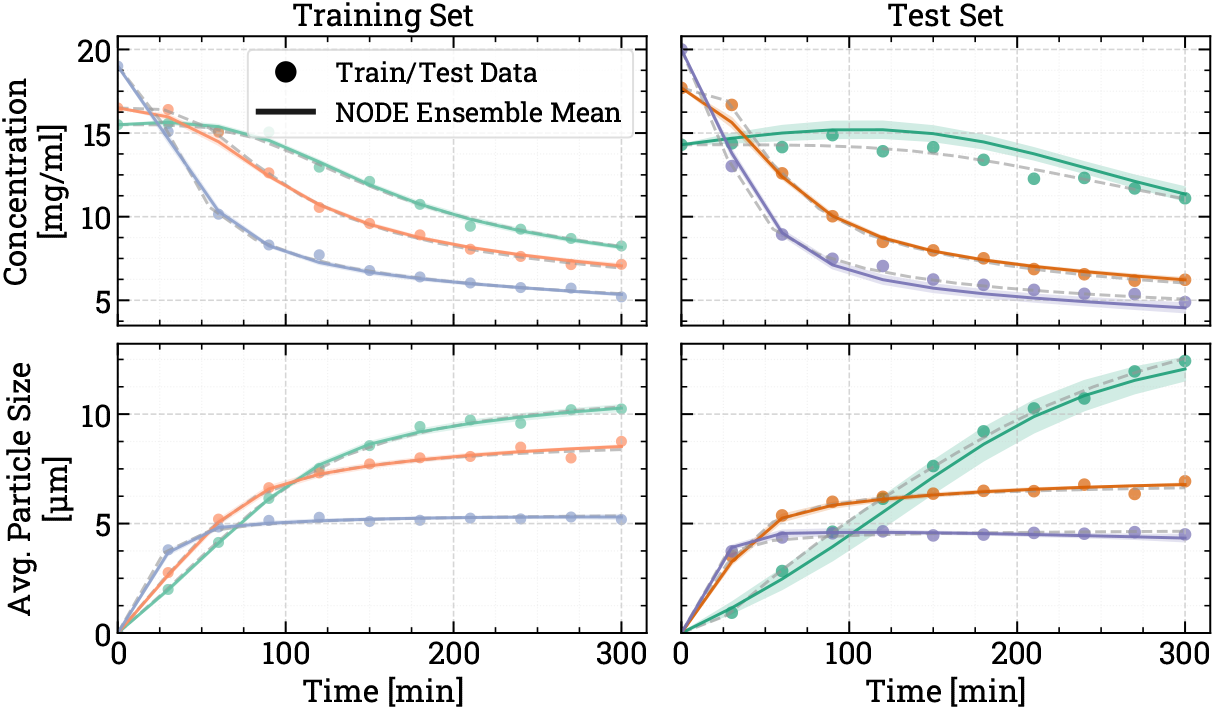
NODE training and testing prediction profiles, trained on the source **S** dataset with 10 measured timepoints and 2% added noise.

**Figure 3:**
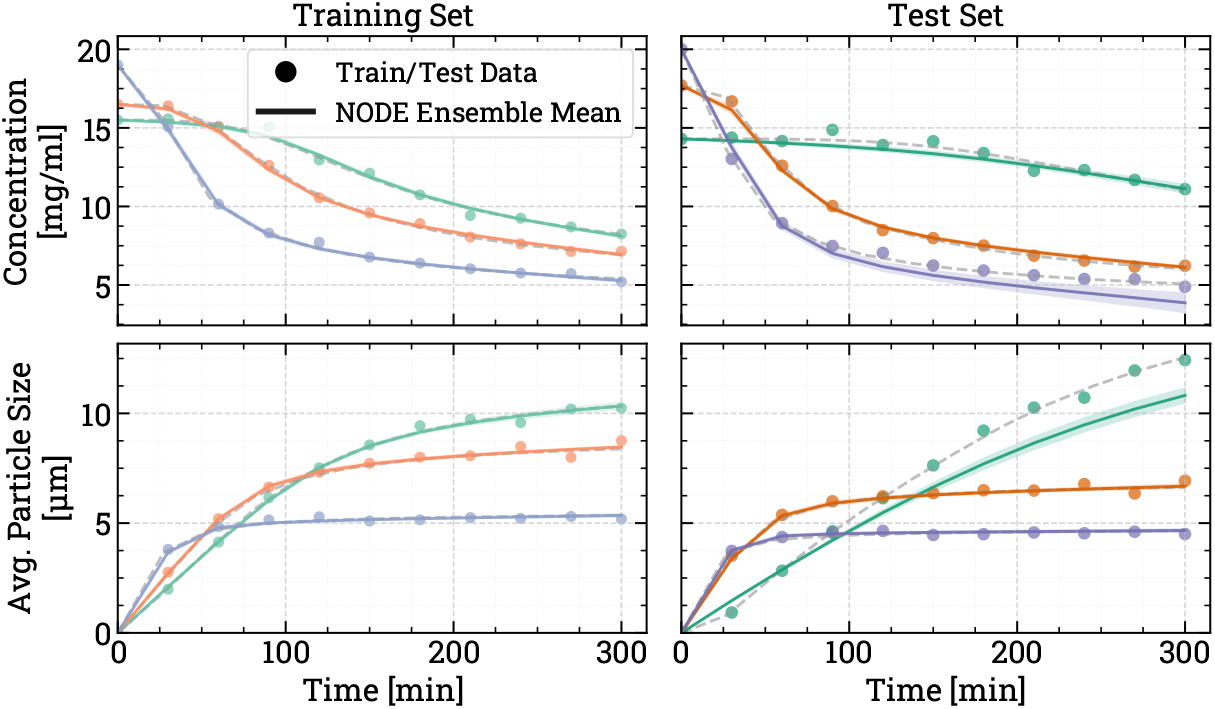
Constrained NODE training and testing prediction profiles, trained on the source **S** dataset with 10 measured timepoints and 2% added noise.

NODEs are also trained on the two-experiment **T**_i_ datasets to serve as baselines for the knowledge-transfer methods assessment, and their predictions are shown in Figures 4, 5 and 6. Generalization to the test data reveals limitations due to the reduced training dataset. For **T**_1_, the NODE fails to predict both solute concentration and average particle size accurately. In contrast, the **T**_2_ and **T**_3_ particle size trajectories are modelled with reasonable accuracy, while concentration predictions remain inaccurate. All scratch-trained models struggle to replicate the longer induction period observed in the test concentration profiles, as the flatter initial segment of the concentration curve is consistently underestimated. This can be attributed to the narrow range of initial conditions in the training data, which fails to expose the NODE to enough variability in induction times, and consequently the NODEs cannot predict it. The scratch-trained 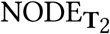 and 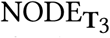 predict accurate particle size profiles.

**Figure 4:**
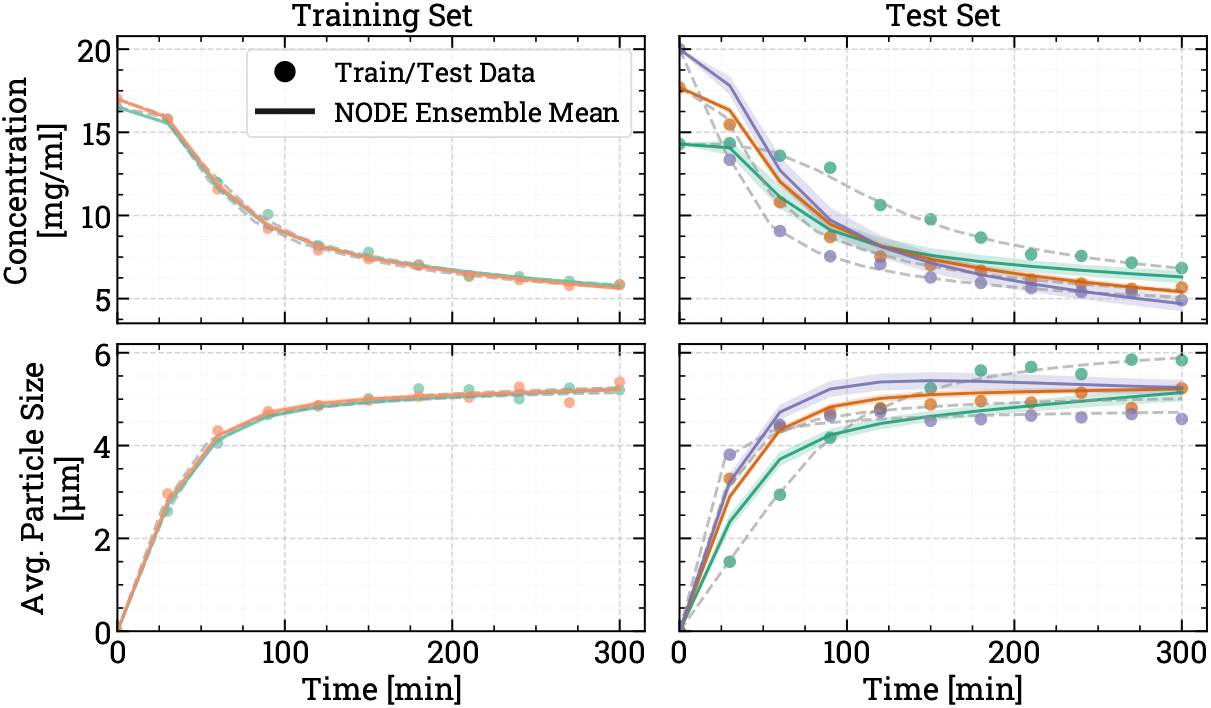
NODE training and testing prediction profiles, scratch-trained on the target **T**_1_ dataset with 10 measured timepoints and 2% added noise.

**Figure 5:**
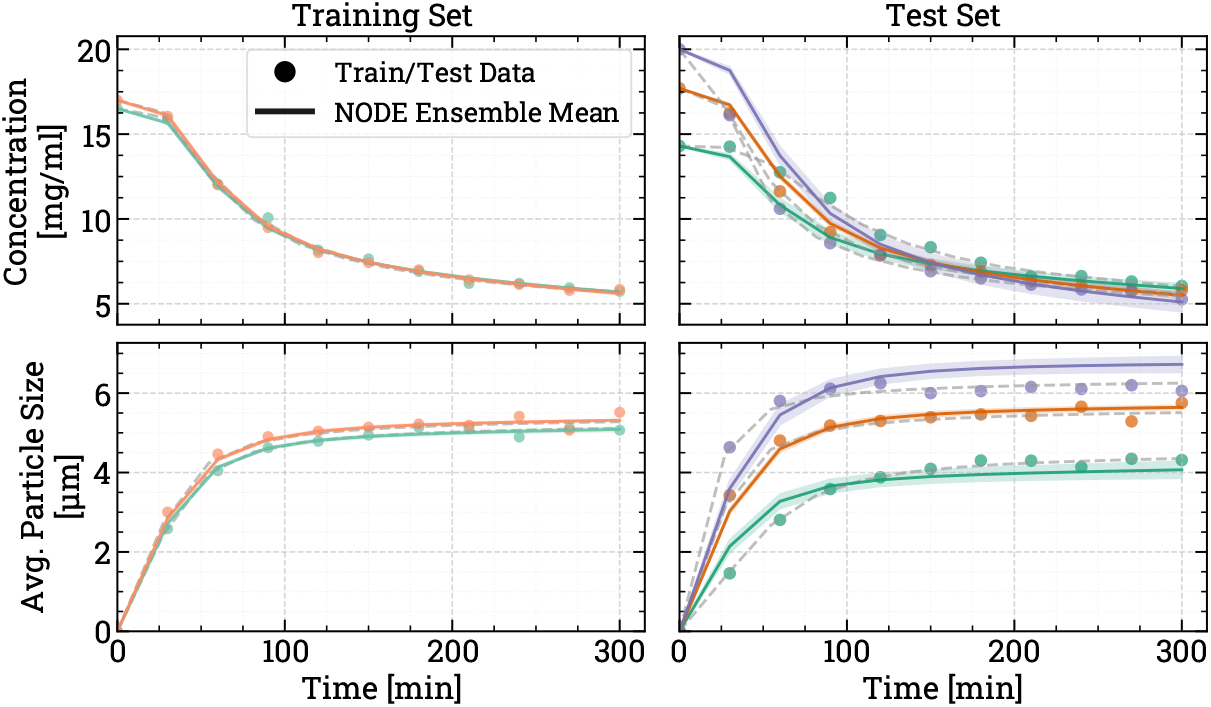
NODE training and testing prediction profiles, scratch-trained on the target **T**_2_ dataset with 10 measured timepoints and 2% added noise.

**Figure 6:**
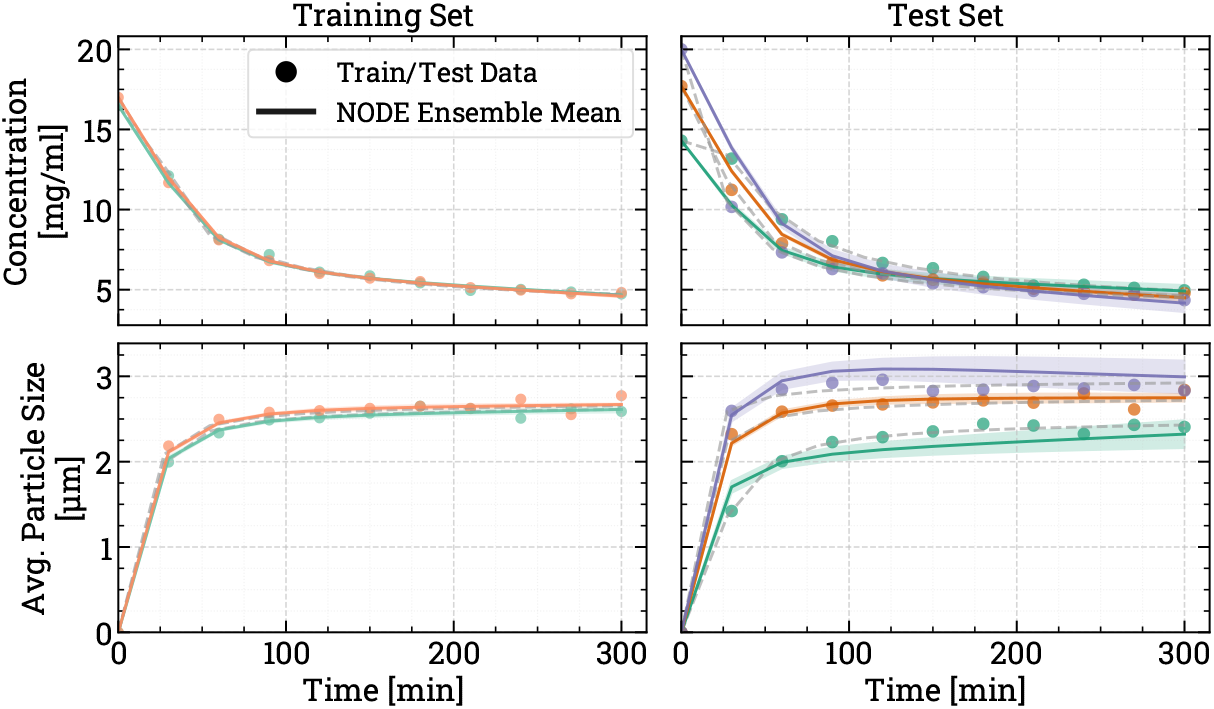
NODE training and testing prediction profiles, scratch-trained on the target **T**_3_ dataset with 10 measured timepoints and 2% added noise.

All profiles show the same trend, with final particle size increasing with the initial solute concentration. This consistent monotonic behaviour likely enables the model to interpolate and extrapolate effectively, even under data scarcity. However, **T**_1_ presents more complex crystallization behaviour, similar to that observed in the source system. Here, the final particle size decreases with increasing initial concentration, mechanistically explained by competing crystallisation kinetics. In the **T**_2_ and **T**_3_ systems, the carboxyl- and butyl-functionalised templates produce very sharp nucleation, which rapidly depletes enough solute to lower the supersaturation and shift the batch into a crystal growth-dominated regime. In contrast, **S** and **T**_1_ exhibit slower nucleation, allowing prolonged competition between nucleation and growth. 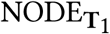 has limited exposure of this competition in its’ training data, and fails to learn and generalise the trend.

### 3.1. Knowledge transfer assessments

#### 3.1.1. Unconstrained NODEs

Given that the unconstrained and constrained source-trained NODE successfully capture both solute concentration and particle size dynamics, various knowledge transfer strategies are employed to adapt the models to the data-scarce target systems, and leverage the learned dynamics from the source domain to improve the predictive accuracy on the **T**_i_ datasets. Each strategy is assessed against each system.

Examining the errors in Figure 7, the embedded NODE offers the best concentration predictions across all three target systems, as it is able to leverage information from all four systems, rather than just **S**-**T**_i_ pairs. The penalisation methods perform better compared to freezing strategies achieving lower or equivalent error in concentration predictions. Average particle size predictions are more accurate when using freezing and penalisation TL methods, compared to finetuning and embedding methods which have the highest MAE. This underperformance likely results from the simplistic embedding mechanism used. The shared linear projection may conflate system-specific *d*_4,3_ kinetics and fail to disentangle critical differences across crystallization regimes. Other implementations of embedded NODEs have adopted richer embedding architectures such as encoder/decoder setups (albeit for different tasks to process modelling), which offer higher expressivity and greater flexibility in learning latent system representations (Alaa et al. 2022; Dang et al. 2023). It’s possible that these more advanced embeddings would be able to better separate the non-linear and system-specific dynamics required for effective generalization across heterogeneous crystallization processes. Overall, freezing- and penalisation-based TL methods all have similar balances of concentration and average particle size errors, and appear to perform equally well.

**Figure 7:**
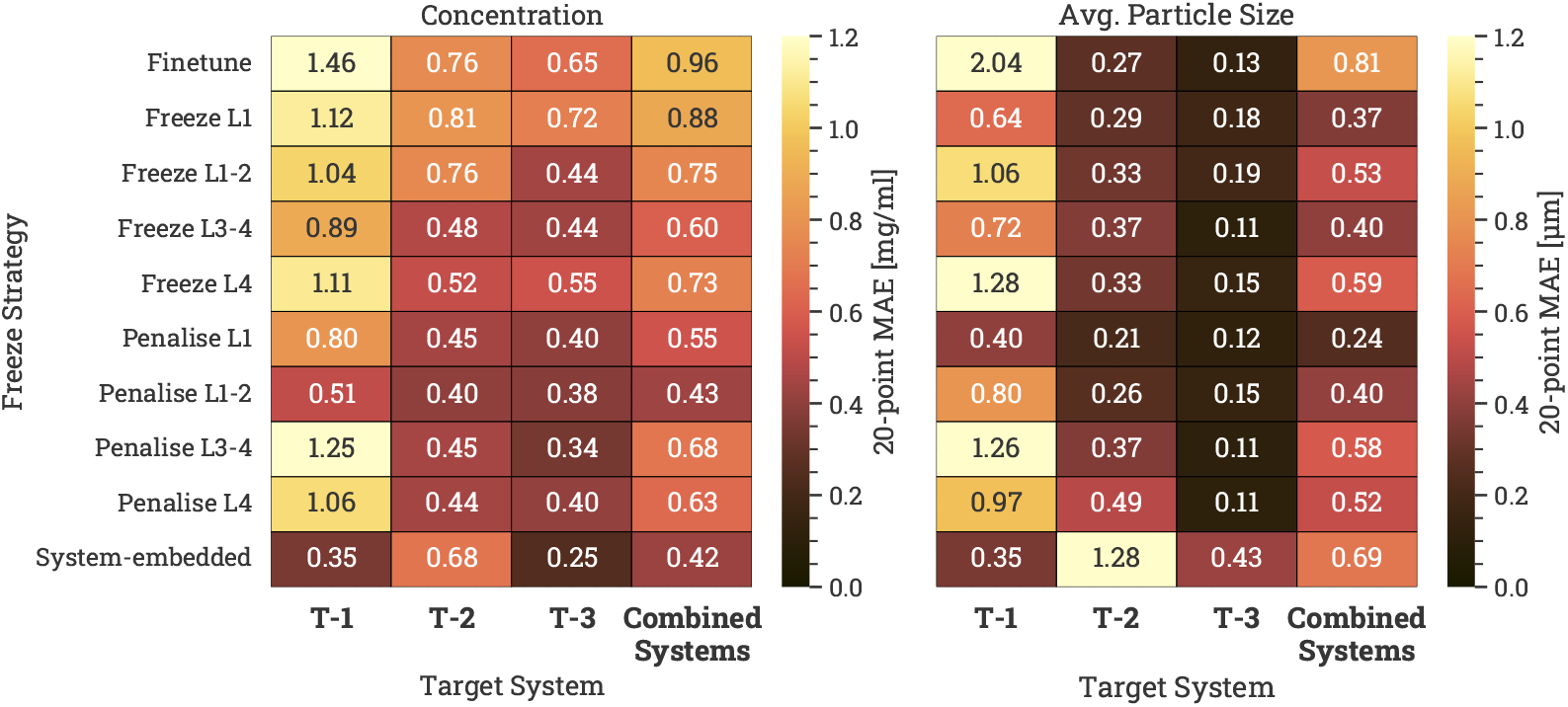
Concentration and *d*_43_ mean absolute error heatmap of the different knowledge transfer methods with unconstrained NODEs. The dataset contains 10 measured timepoints and 2% added noise.

Figures 8, 9 and 10 show predictions from a transferred unconstrained NODE after L4 is frozen. Transfer to **T**_1_ appears to be successful. Compared to the scratch-trained NODE’s predictions in Figure 4, the transferred NODE can accurately capture the crystallisation kinetics; the concentration profile is now reliably modelled, and it includes the previously-missing induction time. The *d*_43_ predictions are also more accurate, with the transferred NODE predicting the competing crystallisation kinetic behaviour and particle size trends of **T**_1_. However, the transferred NODE also predicts a non-monotonic particle size trend for the test experiment at the highest concentration (20 mg/ml), highlighting the poor information diversity in the target training data.

**Figure 8:**
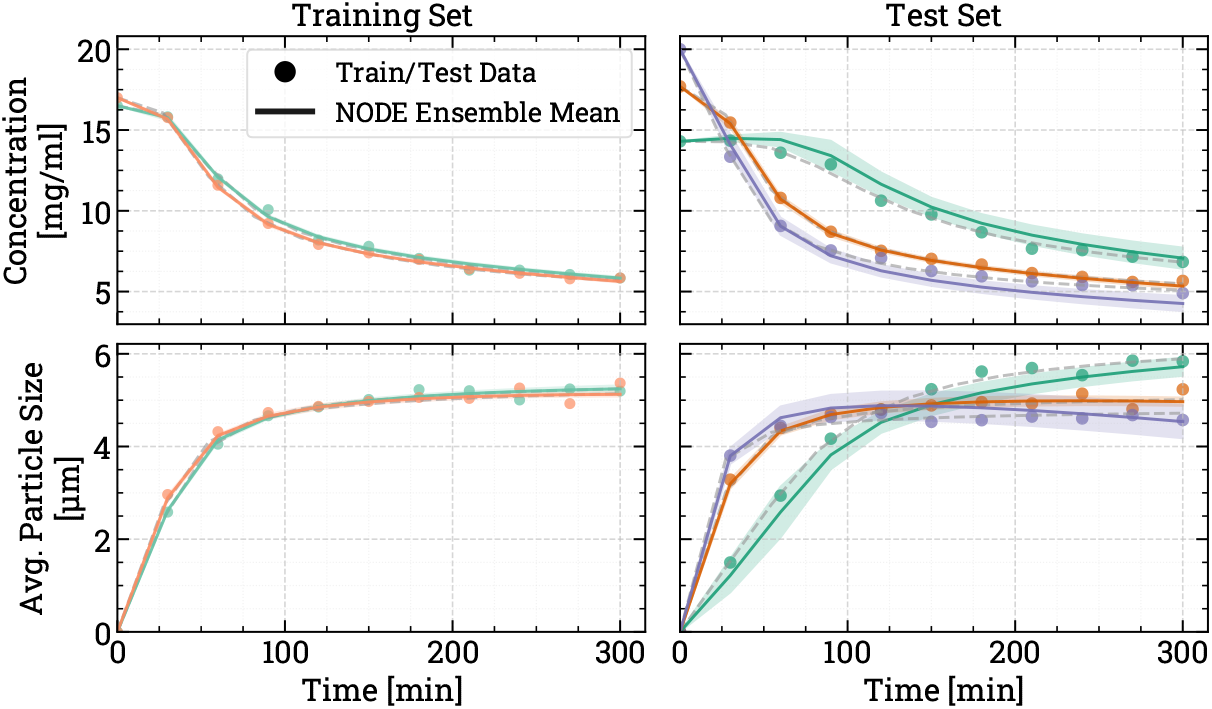
NODE training and testing prediction profiles, trained on the target **T**_1_ dataset with L4 frozen layer, 10 measured timepoints and 2% added noise.

**Figure 9:**
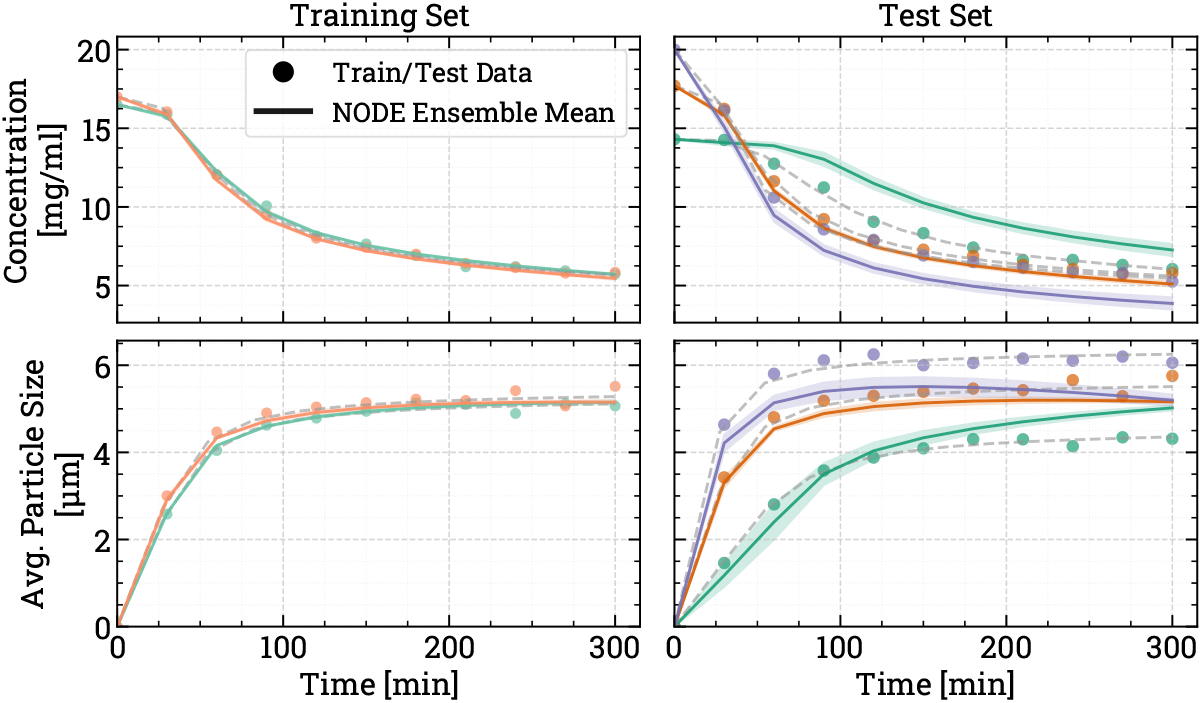
NODE training and testing prediction profiles, trained on the target **T**_2_ dataset with L4 frozen layer, 10 measured timepoints and 2% added noise.

**Figure 10:**
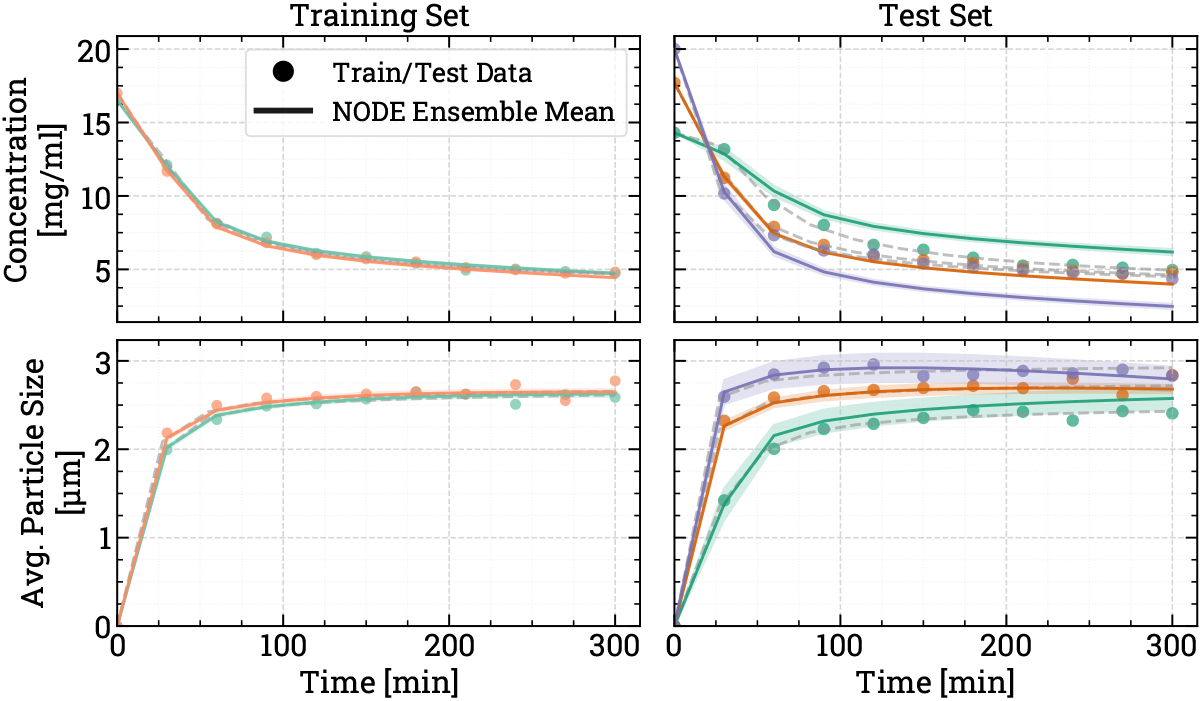
NODE training and testing prediction profiles, trained on the target **T**_3_ dataset with L4 frozen layer, 10 measured timepoints and 2% added noise.

Similar trends are observed for unconstrained NODEs transferred to **T**_2_ and **T**_3_. Both the concentration and average particle size profiles are reliably captured. The 20 mg/ml experiment again has slightly poorer predictions than others. Specifically, the predicted concentration profiles of both systems do not stall and slow down, but are instead approaching complete saturation. The particle size is also non-monotonic, and starts decreasing towards the end of the batch. Since NODEs are extrapolating outside of both the source and target training data, the lack of information on crystallisation kinetics at these conditions leads the NODEs to inaccurate gradient predictions.

#### 3.1.2. Constrained NODEs

Constrained NODEs are also tested for knowledge transfer and compared to their unconstrained counterparts. The concentration and *d*_43_ prediction errors for transferred constrained NODEs (Figure 11) show that constraining the NODE outputs to enforce monotonicity leads to more accurate particle size predictions at the cost of poorer concentration predictions, regardless of the TL method used. The two negative and positive monotonicity constraints for concentration and *d*_43_ respectively, might be affecting the ODE flow learned during NODE training, and worsen the final NODE accuracy. It is however possible that this is also due to the chosen neural network hyperparemeters, since a larger neural network structure or additional augmented dimensions in the input could improve training flexibility and assist NODE training given the output constraints.

**Figure 11:**
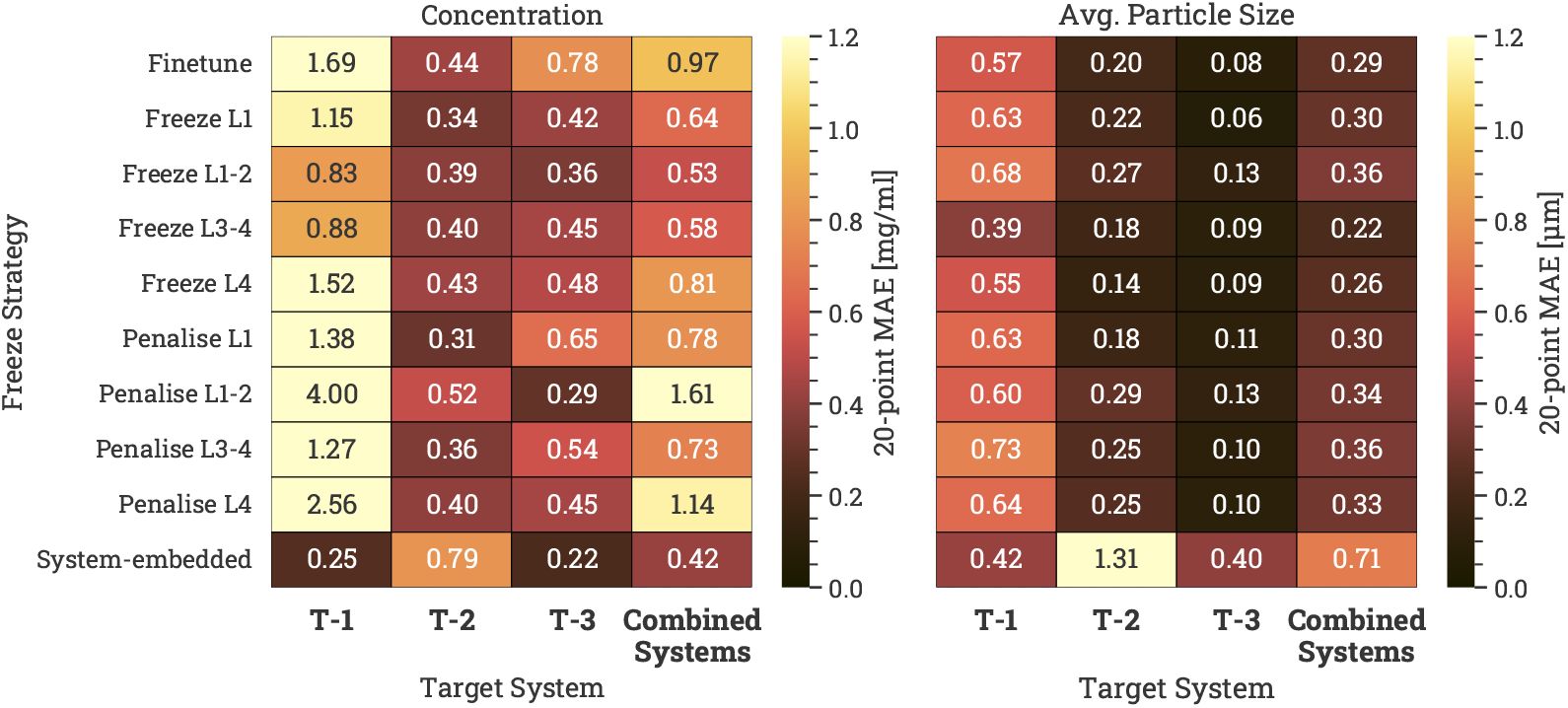
Concentration and *d*_43_ mean absolute error heatmap of the different knowledge transfer methods with constrained NODEs. The dataset contains 10 measured timepoints and 2% added noise.

Prediction profiles of the transferred constrained NODEs with L4 frozen are shown in Figures 12, 13 and 14. The predicted concentration profiles still capture the induction time, but also fail at capturing the concentration’s slower consumption towards the end of the batch. As noted in Figure 11, particle size predictions are instead improved compared to the unconstrained NODEs, as the non-monotonic behaviour observed in Figures 9 and 10 is completely mitigated, and the predictions remain physically consistent.

**Figure 12:**
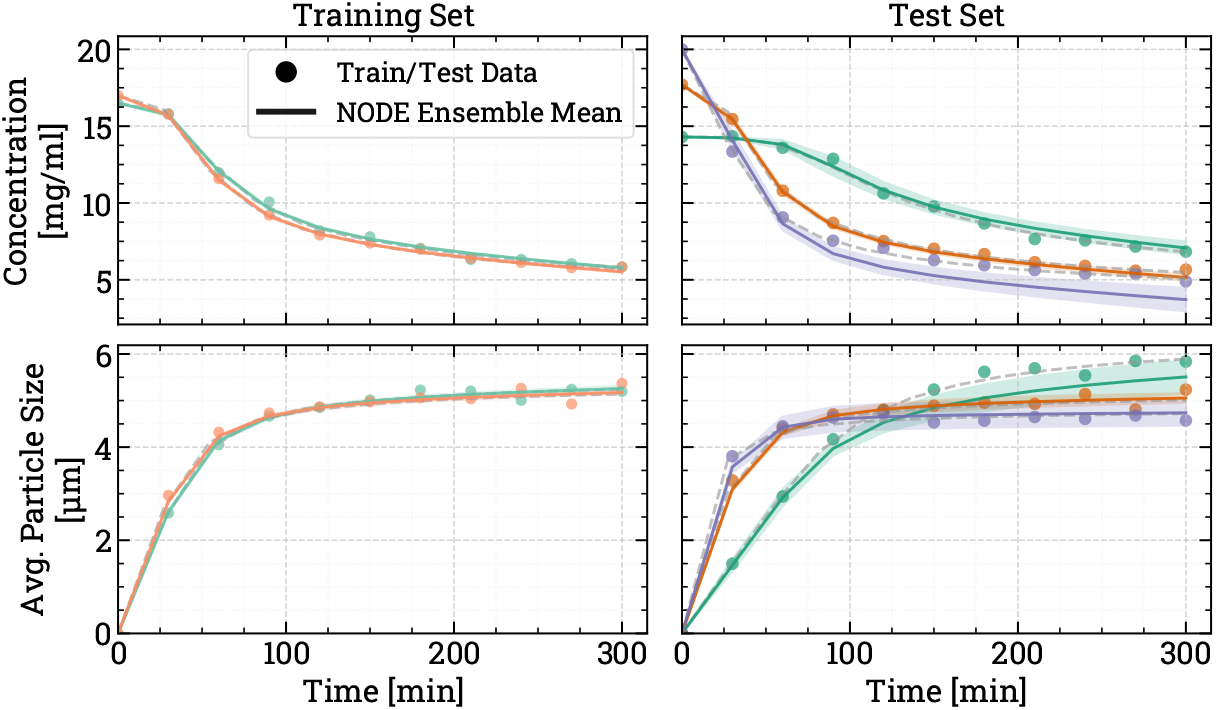
Constrained NODE training and testing prediction profiles, trained on the target **T**_1_ dataset with L4 frozen layer, 10 measured timepoints and 2% added noise.

**Figure 13:**
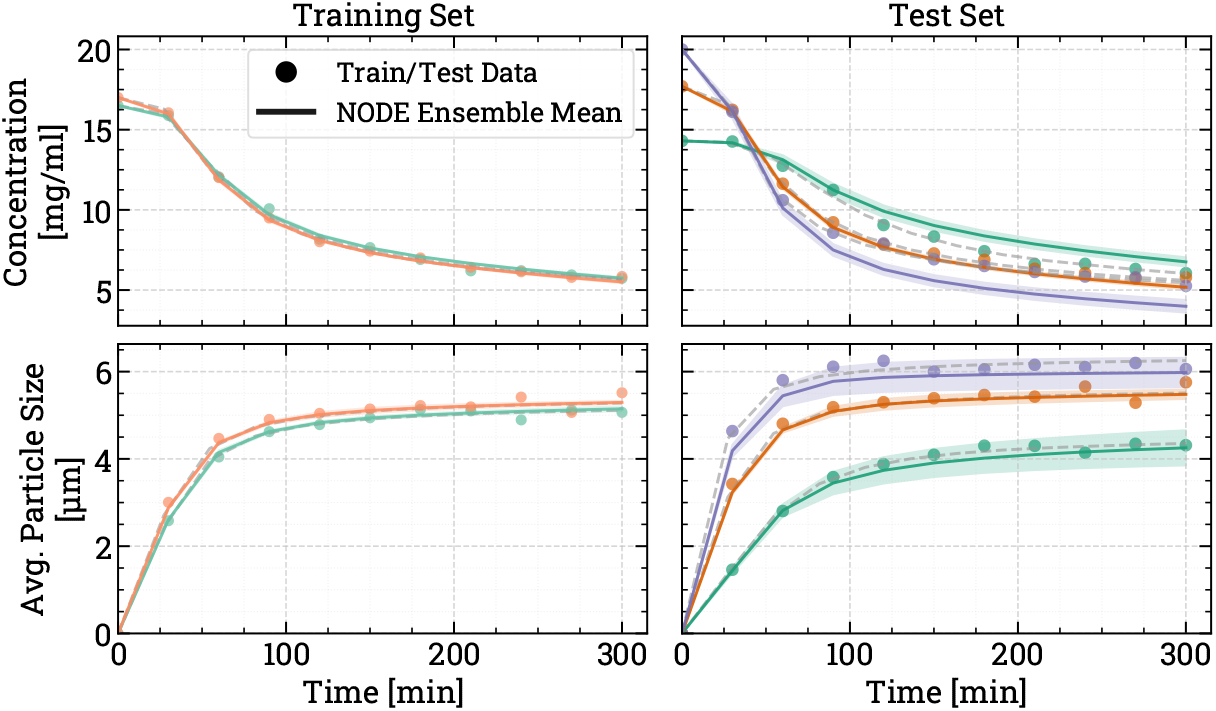
Constrained NODE training and testing prediction profiles, trained on the target **T**_2_ dataset with L4 frozen layer, 10 measured timepoints and 2% added noise.

**Figure 14:**
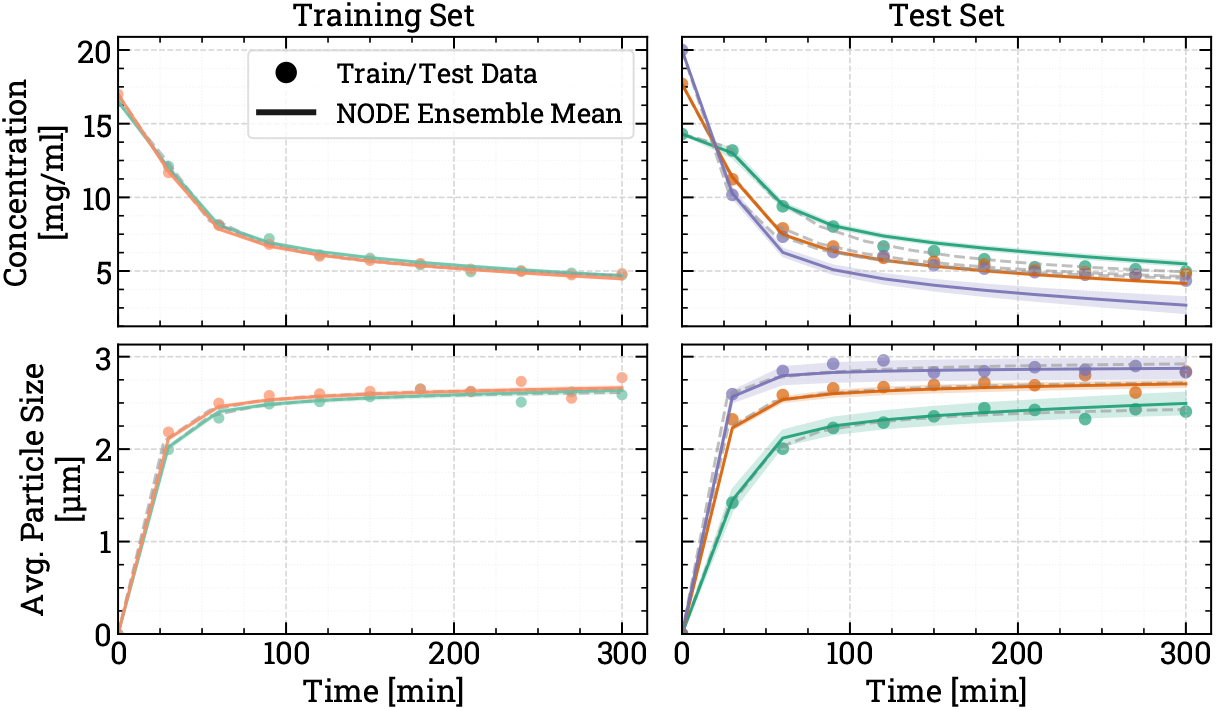
Constrained NODE training and testing prediction profiles, trained on the target **T**_3_ dataset with L4 frozen layer, 10 measured timepoints and 2% added noise.

### 3.2. Robustness study against noise level and number of measured samples

Each knowledge transfer method was also tested for different numbers of measured time points and levels of added gaussian noise, with the aggregated **T**_i_ errors collected in Figures 15 and 16. Higher levels of noise and fewer timepoints in the training datasets are correlated to higher recorded errors in the testing experiments. Sparser datasets negatively affect the knowledge transfer success, as the datasets have fewer informative points used to firstly train a well-predicting source NODE, and secondly transfer the NODE to the target domain and its’ kinetics. The strongest effect however comes from the amount of noise in the dataset. Since the two **T**_i_ training experiments have similar initial conditions and trajectories, high noise in the measurements can mask the more nuanced differences between them, and prevents successful knowledge transfer. Transfer with high-noise data can also make the whole transferred NODE unstable, as evidenced by the high errors when finetuning, or when penalising L1, L3+4 and L4. Embedded NODEs are instead comparatively insensitive to both the amount of noise and the amount of measured samples, again highlighting the advantage of the embedding method to leverage information from all four systems rather than a pair of them.

**Figure 15:**
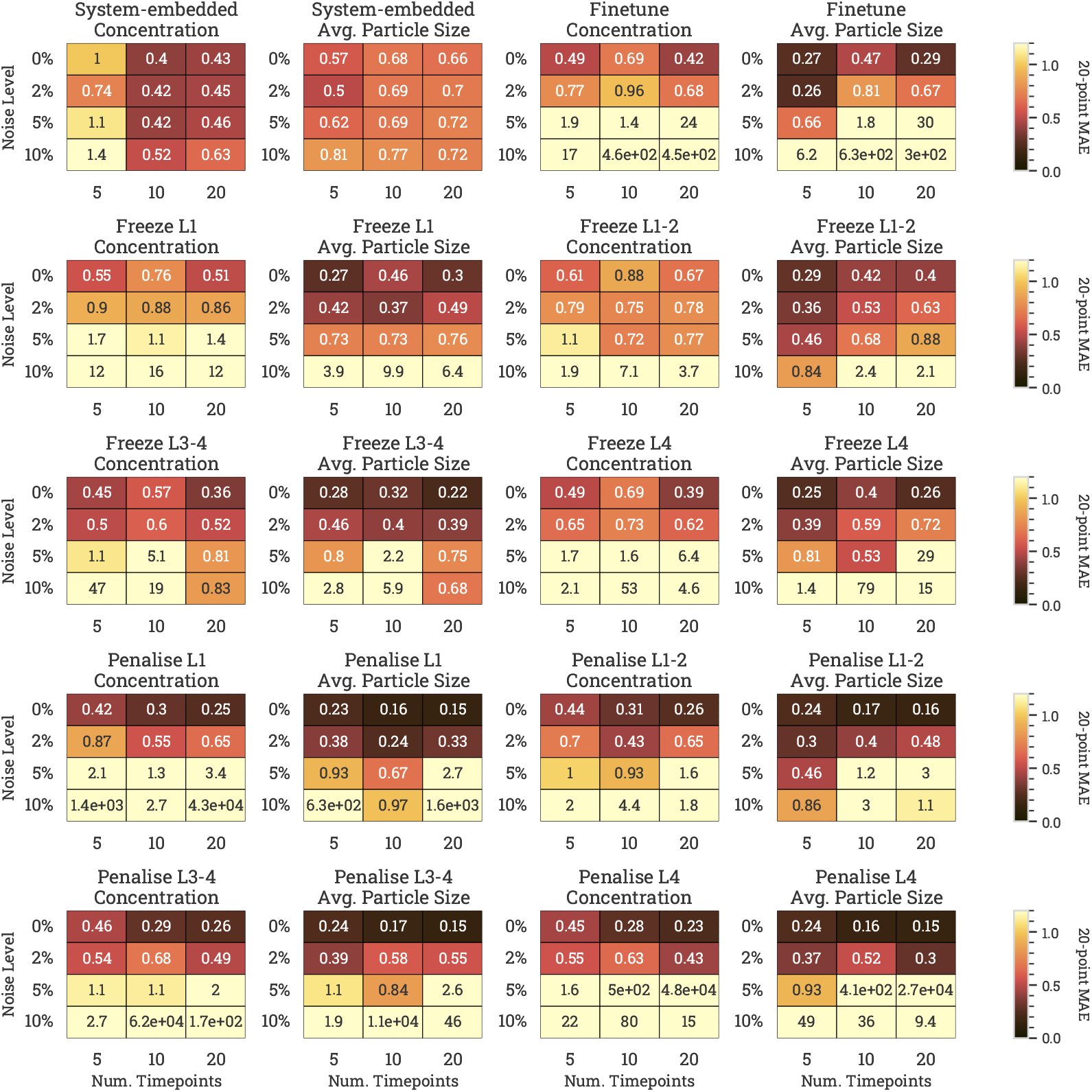
Concentration and *d*_43_ mean absolute error heatmap of the different knowledge transfer methods with unconstrained NODEs for different combinations of added gaussian noise and number of measured timepoints

**Figure 16:**
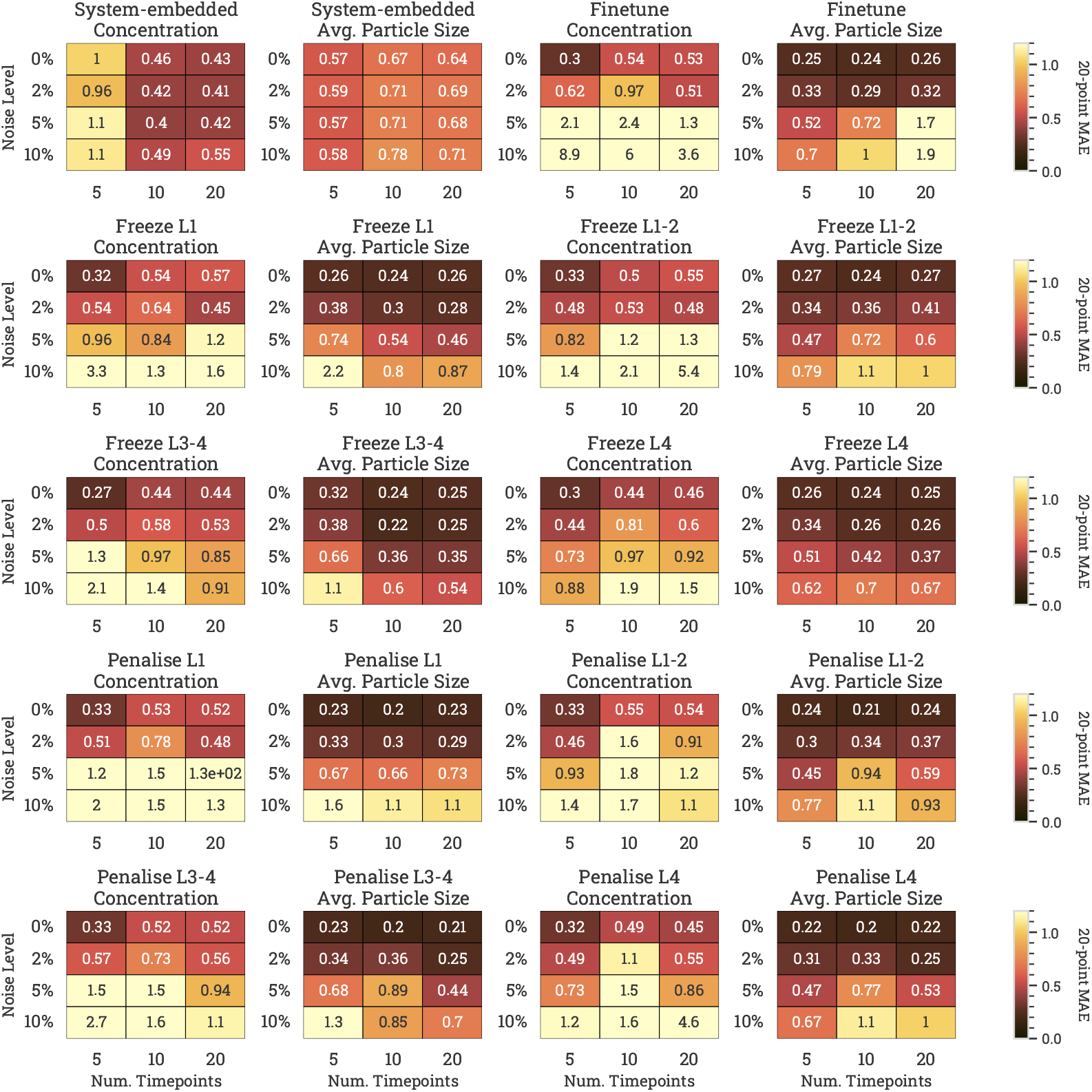
Concentration and *d*_43_ mean absolute error heatmap of the different knowledge transfer methods with constrained NODEs for different combinations of added gaussian noise and number of measured timepoints

Constraining the NODEs has several effects on the success of each TL method. Firstly, constrained NODEs have more stable training during knowledge transfer with noisy data. The post-transfer instability observed for some TL methods with 10% noise in Figure 15 is instead wholly mitigated by the constrained NODE structure. Secondly, the observations made in Sec. 3.1.2 hold for the range of noise and number of samples tested in this robustness study. Constrained NODEs achieve comparable concentration error predictions for the range of noise levels and sample numbers tested, while improving the particle size predictions. By introducing physics-informed constraints to the NODE structure, the resultant post-transfer NODEs are more robust against noisy and scarce datasets, and more accurate compared to an unconstrained NODE.

## 4. Conclusions

Protein crystallisation processes are central to the production of high-purity solid APIs. However, the proteins’ complex dynamics and the limited availability of high-fidelity process data detrimentally affects the predictive performance of both mechanistic and data-driven models. Addressing these challenges is critical for advancing predictive control, optimisation, and process design strategies.

This study demonstrates the potential of Neural ODEs to model complex crystallisation dynamics across distinct systems, even in low-data regimes, and contributes a step toward reliable, data-efficient neural modelling of crystallisation and more broadly of scientific dynamical systems. Unconstrained and physics-informed NODEs are formulated by enforcing monotonicity constraints of the two model outputs. The NODE trained on the source dataset successfully learns the underlying concentration and particle size dynamics, and serves as a robust foundation for knowledge transfer.

Among the transfer learning strategies explored, layer freezing and deviation penalty consistently achieved reliable knowledge transfer, offering balanced performance across concentration and particle size predictions. Embedding methods, which can leverage information from all four systems simultaneously, is strongest when the dataset is noisy or has few measured points. Finally, the comparison between unconstrained and constrained NODEs highlights an important advantage. The unconstrained NODE’s performance deteriorated under noisy or sparse conditions, occasionally yielding unphysical behaviours, while constrained NODEs were systematically more robust in the presence of high noise and limited measurements, maintaining physical consistency and stabilising post-transfer training. This suggests that physics-informed constraints, when combined with carefully selected transfer learning methods such as layer freezing or deviation penalty, provide a powerful framework for enabling reliable knowledge transfer of NODEs in low-data crystallisation domains.

## Supporting information

Crystallisation ODE Model

