## Supplementary material for "Transfer Learning Assessment of Data-driven Crystallisation Processes via Constrained Neural Ordinary Differential Equations": Crystallisation ODE Model

### Appendix A. Crystallisation ODE model

An *in-silico* crystallisation model is used to generate the data necessary for this work. Eq. A.1 is the population balance equation (PBE) for a batch crystallisation process with negligible aggregation and breakage. Given it is a hyperbolic partial differential equation, specific solution methods must be used to solve the PBE. The PBE is solved using the Method of Moments (Eqs. A.2-A.7), which returns moments of the crystal size distribution rather than a solution of the crystal number density $n$ (Randolph et al. 1988).

$$\frac{\partial \overset{\text{Crystal number density}}{n}}{\partial t} + \frac{\partial (Gn)}{\partial L} = B_0 \delta(\overset{\text{Crystal length}}{L} - L_{\text{min}}) \tag{A.1}$$

$$\frac{d\mu_0}{dt} = \overset{\text{Nucleation rate}}{B_0} \tag{A.2}$$

$$\frac{d\mu_1}{dt} = G\mu_0 \tag{A.3}$$

$$\frac{d\mu_2}{dt} = 2G\mu_1 \tag{A.4}$$

$$\frac{d\mu_3}{dt} = 3G\mu_2 \tag{A.5}$$

$$\frac{d\mu_4}{dt} = 4G\mu_3 \tag{A.6}$$

$$\frac{dc}{dt} = -3 \underset{\text{Volumetric shape factor}}{k_v} \underset{\text{Crystal density}}{\rho_c} \underset{\text{Growth rate}}{G} \underset{\text{Moment of the distribution}}{\mu_2} \tag{A.7}$$

The model equations are collected in Eqs. A.2-A.7. Eq. A.7 is the mass balance of the crystalliser, and is used to calculate solute consumption during the batch. The supersaturation $S$ (Eq. A.8), defined as the ratio between solute concentration and saturation concentration, is the thermodynamic driving force of the process. Classical Nucleation Theory and an empirical power-law growth equation are used to respectively model the nucleation rate $B_0$ and growth rate $G$. Finally, the volume-weighted average particle size $d_{43}$ is calculated using Eq. A.11. The relevant kinetic, physical and thermodynamic parameters are collected in Tables A.1 and A.2. The saturation concentration $c_{\text{sat.}}$ is calculated from correlations provided by Cacioppo et al. (1991). The density of a tetragonal lysozyme crystal $\rho_c$ is given by Leung et al. (1999). Lysozyme's molecular volume $v_0$ is given by Nadarajah et al. (1996).

Solute concentration

$$S = \frac{c}{c_{\text{sat.}}} \tag{A.8}$$

$$B_0 = A_j S \exp\left[\frac{-16\pi\gamma^3 v_0^2}{3k_B^3 T^3 \ln^2 S}\right] \tag{A.9}$$

$$G = A_g (S-1)^g \tag{A.10}$$

$$d_{43}(t) = \frac{\mu_4}{\mu_3} \tag{A.11}$$

| **System** | **Nucleation Parameters** | |
|---|---|---|
| | $A_j$ [log # $m^{-3}s^{-1}$] | $\gamma$ [mJ/$m^{-2}$] |
| Unseeded **S** | 39.8 | 0.675 |
| Hydroxyl $\mathbf{T}_1$ | 29.9 | 0.49 |
| Carboxyl $\mathbf{T}_2$ | 22.7 | 0.14 |
| Butyl $\mathbf{T}_3$ | 27.4 | 0.32 |
| | **Growth Parameters** | |
| – | $A_g$ [nm $min^{-1}$] | $g$ [–] |
| – | 0.37 | 3.3 |

Table A.1: Nucleation and growth kinetic parameters used to generate *in-silico* data

| Parameter | Name | Value |
|---|---|---|
| $c_{sat.}$ | Saturation concentration | 2.47 [mg ml$^{-1}$] |
| $k_v$ | Volumetric shape factor | 0.81 [-] |
| $\rho_c$ | Crystal Density | 1240 [kg m$^{-3}$] |
| $v_0$ | Molecular volume | 2.97 $\times 10^{-26}$ [m$^{-3}$] |
| $k_B$ | Boltzmann constant | 1.38 $\times 10^{-23}$ [m$^2$ kg s$^{-2}$ K$^{-1}$] |

Table A.2: Physical and thermodynamic parameters used to generate *in-silico* data

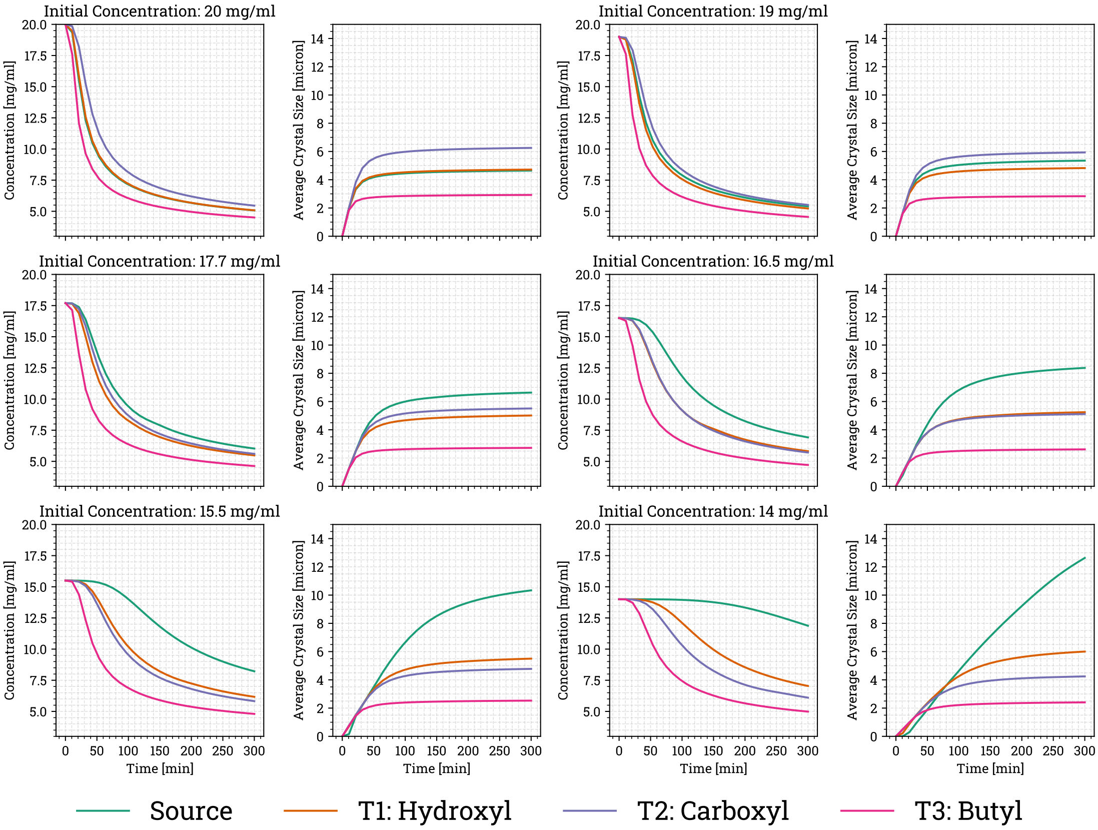


Figure A.1: Simulated solute concentration and average particle size trajectories
